# Comparative genomic analysis of *Mycobacterium tuberculosis* reveals evolution and genomic instability within Uganda I sub-lineage

**DOI:** 10.1101/2020.10.24.353425

**Authors:** Stephen Kanyerezi, Patricia Nabisubi

**Affiliations:** The African Centers of Excellence in Bioinformatics and Data Intensive Sciences, the Infectious Diseases Institute, College of Health Science, Makerere University P.O Box 22418 Kampala, Uganda

## Abstract

**Introduction:** Tuberculosis (TB) is the leading cause of morbidity and mortality globally, responsible for an estimated annual 10.0 million new cases and 1.3 million deaths among infectious diseases with Africa contributing a quarter of these cases in 2019. Classification of *Mycobacterium tuberculosis* (MTB) strains is important in understanding their geographical predominance and pathogenicity. Different studies have gone ahead to classify MTB using different methods. Some of these include; RFLP, spoligotyping, MIRU-VNTR and SNP set based phylogeny. The SNP set based classification has been found to be in concordance with the region of difference (RD) analysis of MTB complex classification system. In Uganda, the most common cause of pulmonary tuberculosis (PTB) is Uganda genotype of MTB and accounts for up to 70 % of isolates.

**Methods:** Sequenced MTB genome samples were retrieved from NCBI and others from local sequencing projects. The genomes were subjected to snippy (a rapid haploid variant calling and core genome alignment) to call variants and annotate them. Outputs from snippy were used to classify the isolates into Uganda genotypes and Non Ugandan genotypes based on 62 SNP set. The Ugandan genotype isolates were later subjected to 413 SNP set and then to a pan genome wide association analysis.

**Results:** 6 Uganda genotype isolates were found not to classify as either Uganda I or II genotypes based on the 62 SNP set. Using the 413 SNP set, the 6 Uganda genotype isolates were found to have only one SNP out of the 7 SNPs that classify the Uganda I genotypes. They were also found to have both missense and frameshift mutations within the *ctpH* gene whereas the rest of Uganda I that had a mutation within this gene, was a missense.

**Conclusion:** Among the Uganda genotypes genomes, Uganda I genomes are unstable. We used publicly available datasets to perform analysis like mapping, variant calling, mixed infection, pan-genome analysis to investigate and compare evolution of the Ugandan genotype.

## Background

Tuberculosis (TB) is the leading cause of morbidity and mortality globally, responsible for an estimated annual 10.0 million new cases and 1.3 million deaths among infectious diseases with Africa contributing a quarter of these cases by 2019(*WHO | Global Tuberculosis Report 2020*). Uganda was ranked as the 16th among the 22 countries with the heaviest burden of TB with an incidence of 299 and a mortality of 84 cases per 100,000 per year(*WHO | Global Tuberculosis Report 2020*,). Tuberculosis is caused by *Mycobacterium tuberculosis* pathogen of humans and animals. It is made of a complex called the *Mycobacterium tuberculosis* Complex (MTBC)(Riojas et al., 2018), which comprises of different lineages, some referred to as *Mycobacterium tuberculosis sensu stricto* (Lineage 1-4, 7) and others as *Mycobacterium africanum* (Lineage 5 and 6), a recently discovered Lineage 8(Ngabonziza et al., 2020) in Ugandan and Rwandan tuberculosis patients as well as lineage 9(Mireia Coscolla et al., 2020) in East Africa. Furthermore, MTBC also consists of animal - associated ecotypes which include; *M. bovis, M. pinnipedii*, or *M. microti* among others. Human lineages are geographically widespread and others are restricted like the case for Lineage 7 that is limited to the Horn of Africa, and L5 and L6 that are mainly found in West Africa (Ngabonziza et al., 2020), (Brites et al., 2018), (S, 2018), (Gagneux et al., 2006), (Firdessa et al., 2013). Lineage 2 origins are in East Asia(Merker et al., 2015), (Luo et al., 2015) and has expanded to some parts of the world(Cowley et al., 2008). Lineage 4 is globally distributed contrary to the other main human-adapted MTBC lineages, but the reasons for this are still unknown,(Demay et al., 2012) though its strains are geographically restricted like the Ugandan *M. tuberculosis* genotype restricted to Uganda population.

TB clinically evinces in two forms, pulmonary TB and extrapulmonary TB. Most people die from extrapulmonary TB (EPTB) despite pulmonary TB (PTB) being the major type of TB(Sharma & Mohan, 2004), (Golden & Vikram, 2005). In 2015, the Ugandan *M. tuberculosis* genotype was accounted the major cause of pulmonary tuberculosis in Uganda although it has subdue to not causing pulmonary cavitation. The polymerase chain reaction method empowered the discovery of the Uganda genotype(*Threonine 80 Alanine at position 7539*) marker. Uganda genotype MTB being highly prevalent in the Ugandan population, called for studies to further characterize it basing on the spoligotyping method. It was found out that the most prevalent Uganda genotype sublineage lacked hybridisation to spacer 40 and 43 and was named Uganda I. Whereas the least prevalent which lacked only spacer 40 was named Uganda II. (Wamala et al., 2015). Uganda I is also known as lineage 4.6.1.1 whereas Uganda II is lineage 4.6.1.2. (Malm et al., 2017).

There is an urgent need for eradication of tuberculosis which requires a deeper understanding of the biology of MTB. Comparative genomics has exploited the evolution of *Mycobacterium tuberculosis* into lineages, sub lineages and their origins thus empowering tuberculosis epidemiological studies. Work from various studies suggests a plausible role for strain diversity in human tuberculosis within the MTBC. (Mireilla Coscolla & Gagneux, 2010). Many methods for discriminating the *Mycobacterium tuberculosis* complex into it’s distinct lineages have been developed. These include; *IS6110* restriction fragment length polymorphism (RFLP), spacer oligotyping (Spoligotyping), and mycobacterial interspersed repeat units - variable number of tandem repeats (MIRU-VNTR). (Desikan & Narayanan, 2015). To classify MTBC, regions of difference are usually used and are commonly regarded as the gold standard markers. (Faksri et al., 2016). Coll et al., 2014, suggests that the SNP-based phylogeny is congruent with the gold-standard regions of difference (RD) classification system. Comas et al., 2009, Homolka et al., 2012, Feuerriegel et al., 2014, Abadia et al., 2010, Stucki et al., 2012 and Coll et al., 2014 have all proposed the use of several sets of single-nucleotide polymorphisms to classify the MTBC up to the sublineage level. Among these sets, most tools developed to classify the MTBC base on the 62 SNPs randomly proposed by Coll et al., 2014. Some of those tools include snpit (Fowler, 2018/2020), Biohansel (*Phac-Nml/Biohansel*, 2017/2020) and TB-Profiler (Phelan, 2016/2020). Our study focuses on the use of SNPs proposed by Coll et al., 2014 to classify the Uganda genotype MTB.

## Results

### Population structure

A total of 1556 *M. tuberculosis* genomes were analysed using snippy that called variants and annotated them for the downstream analysis. From the snippy results, customized scripts were used to classify the isolates into Ugandan and non Ugandan genotypes based on the 62 synonymous SNP set proposed by Coll et al., 2014. Of the total isolates, 1231 were non Ugandan genotypes and 325 Ugandan genotypes. Among the Ugandan genotypes, 188, were found to be Uganda I, 144 Uganda II and 23 could neither be classified into the two sublineages. To explore the evolutionary relationship of these 23 genomes in the context of Ugandan genotype *M. tuberculosis* sublineage diversity, more analysis was executed to evince a novel or an evolutioned Uganda genotype.

### Mixed infection analysis

Using samtools, the genomes were subjected to a coverage quality check to find out whether they had been efficiently sequenced at the positions that classify Uganda I and Uganda II genotypes. 9 of them had a poor coverage at the position that is indicative of Uganda II genotype and these were excluded in the downstream analysis. The remaining 14 genomes were checked for the possibility of mixed infections using TB-profiler. 8 of these were mixed infections and were neither Uganda 1 or 2 nor had mixed infections. The study proceeded with the 6 genomes associated with lineage 4.6.1 signifying that they belonged to the Uganda genotype but its sub lineage was not known well as Uganda I and II were lineage 4.6.1.1 and lineage 4.6.1.2 respectively.

### Phylogenetic construction

Phylogenetic analysis was done to investigate the relationship of the 6 genomes with the Uganda I and II sub-lineages. Each ID was relabelled by adding ug1, ug2 and notag suffixes in correspondence to their identity as Uganda I, II and neither I nor II respectively. Multiple core SNP site alignment was performed using snippy-core. To infer phylogeny, ete3 build was used to give a maximum likelihood phylogenetic tree. The 6 notag genomes were shown to cluster among the Uganda I sub lineage as shown in fig1. This triggered more analysis to investigate their difference and similarity with Uganda I.

**Fig 1:**
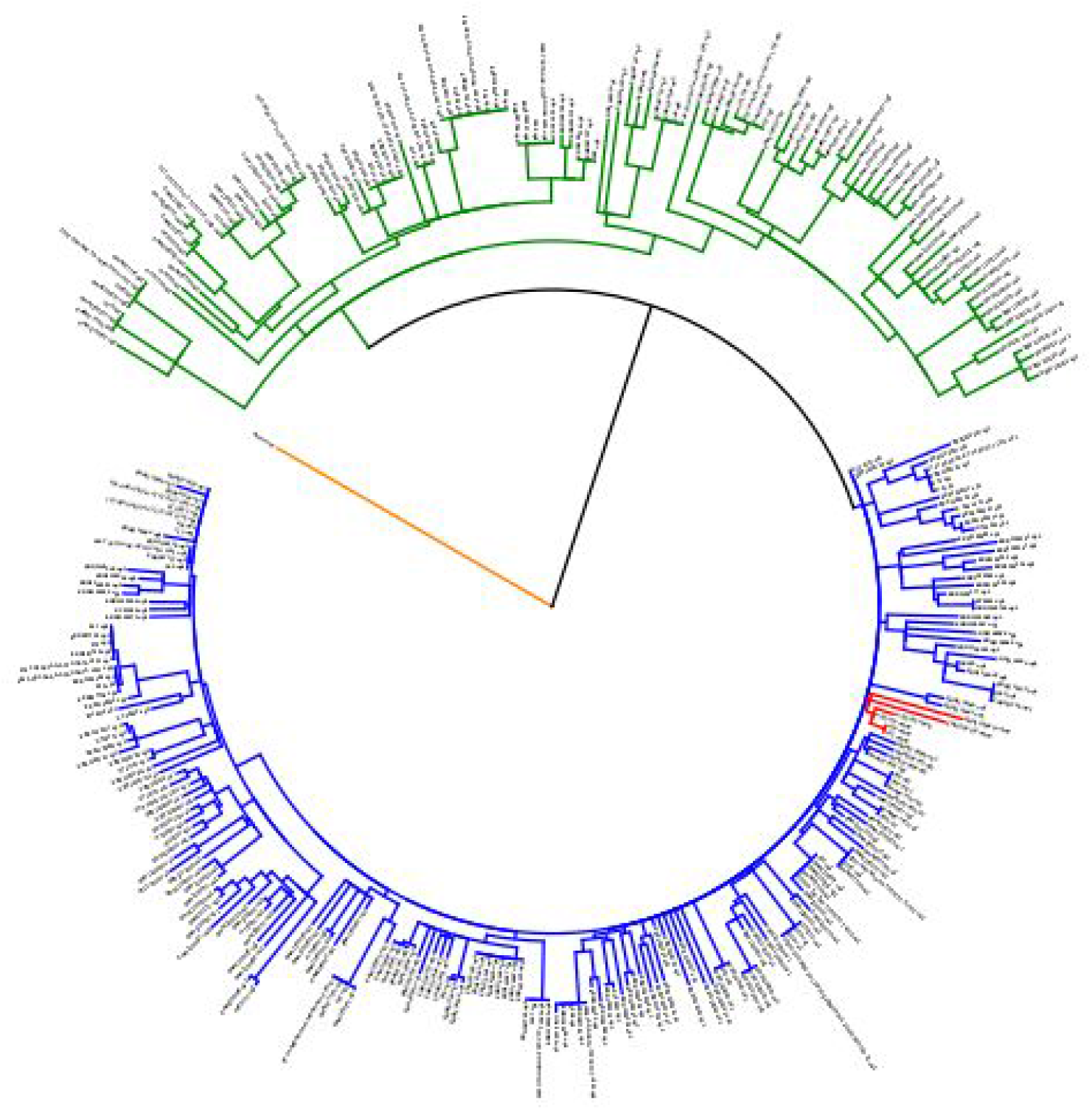
the green clade is for Uganda II, blue for Uganda I and red for notags clade as well as orange is for the reference genome.

### Usage of another SNP set

With the 62 SNP set, the 6 notag isolates were distinguished from the Uganda I isolates yet the phylogenetic tree showed the 6 notag isolates clustering with the Uganda I. Using customized scripts, all the Uganda genotypes were subjected to the 413 synonymous and non synonymous (Coll et al., 2014) SNP set from which the 62 SNP set had been randomly selected. Among the 413 SNPs, 4, 7, 6 SNPs were specific to Uganda, Uganda I and Uganda II genotypes respectively. All the 6 notag isolates had only one SNP (Rv3610c at position 4052608) that classified them as Uganda I. Among the 188 Uganda I isolates, 15 of them lacked both Rv1730c at position 1955910 and Rv3918c at position 4406749 SNPs while the rest had all the 7 SNPs. All Uganda II had all the 6 SNPs. Among all the isolates, only one Uganda I isolate had no one of the SNPs (Rv3800c at position 42602680) classifying Uganda genotypes. To distinguish the 6 notag isolates from Uganda I isolates, a pan-genome analysis was performed.

### Pan-genome analysis

Denovo based assembly of the 325 Uganda genotype isolates was performed using SPAdes genome assembler. The resulting scaffold fasta files containing the assemblies were checked for quality using QUAST. All the assemblies passed the quality check and got annotated by prokka. Roary calculated the pan genome giving an output of the gene presence absence file which, together with the traits file were used by scoary to perform a pan genome wide analysis. There were three traits in the traits file. These included UG1, UG2 and NOTAG for Uganda I, II and the 6 genomes that didn’t classify either as I or II respectively. For each trait, a single csv file consisting of genes that were found to be associated with the respective trait at p < 0.05 was produced. From the NOTAG csv file, all the 6 notag Uganda genomes and 4 other genomes were found to be associated with the “ctpH” gene with a sensitivity of 100.0 and a specificity of 98.39. Using customized scripts, the ctpH gene was further analysed to investigate the type of variants it was associated with. The 6 genomes had the *ctpH* gene associated with the frameshift mutation at position 513274 and a missense mutation at position 513257. Most of the Uganda I and a few Uganda II genomes had the missense mutation at position 513257 but lacked the frameshift mutation at position 513274. The missense mutation was as a result of a single nucleotide change of thymine to cytosine in the reverse strand of the coding sequence leading to an amino acid change of methionine to valine at position 689. The frameshift variant is a result of adenine deletion in the coding sequence of the reverse strand leading to a frameshift of tryptophan.

### Antimycobacterial resistance determining virulence

Since Kasule et al., 2016, Asiimwe et al., 2008 and Lukoye et al., 2014, have all found out the Uganda genotype MTB strains being susceptible to anti-TB drugs, we went on to find out whether the 6 notag isolates do conform to this finding. Mykrobe was used to test for their susceptibility to amikacin, capreomycin, ciprofloxacin, ethambutol, isoniazid, kanamycin, moxifloxacin, ofloxacin, pyrazinamide, rifampicin and streptomycin. All isolates apart from ERR1199118_notag, were susceptible to all the drugs. ERR1199118_notag was susceptible to all the drugs except isoniazid.

## Discussion

This study was designed to understand the evolutionary relationships between the Uganda genotype sub-lineages using whole genome sequenced MTB strains. Among the Uganda genotype isolates, 6 didn’t classify as either Uganda I or II based on the 62 SNP set. This shows that there could have been some missing data during the benchmarking of the SNP set. Using the 413 SNP set, the 6 notag isolates did classify as Uganda I but with only one SNP out of the 7 SNPs that classify Uganda I. Among those that had earlier classified as Uganda I, 15 of them lacked 2 of the SNPs (Rv1730c at position 1955910 and Rv3918c at position 4406749) that are part of those that do classify Uganda I and one of them lacked one of the snp (Rv3800c at position 4260268) markers for Uganda genotype when subjected to the 413 SNP set. For the Uganda II genotype isolates, all of them had the 6 SNPs when subjected to the 413 SNP set. This clearly shows instability within the Uganda I genotype genome.

The 6 genomes were also found to be associated with a frameshift mutation at position 513274 and a missense mutation at position 513257 in the *ctpH* gene though Uganda I with this gene only has the missense mutation. The *ctpH* gene is responsible for production of the P-Type ATPase enzyme for transportation of alkaline metals and also balances the ion influx within the MTB. *MTB* has 11 P-type ATPases (CtpA, CtpB, CtpC, CtpD, CtpE, CtpF, CtpG, CtpH, CtpI, CtpJ, and CtpV). Seven of these enzymes have been identified as possible carriers of heavy metal cations, suggesting their possible role in the intra phagosomal survival of *Mtb*. (Novoa-Aponte et al., 2012), (Argüello et al., 2011). Metals, such as iron, magnesium, cobalt, copper, manganese, and zinc, are essential for all forms of life, as they act as members of prosthetic groups or as cofactors of many enzymes. In general, microorganisms only need traces of these micronutrients for proper cell function; indeed, excessive accumulation is toxic.(Forrellad et al., 2013) An overabundance of metals that can block enzymatic functional groups, as it displaces essential metal ions, and modify the active conformations of biomolecules (Rathnayake et al., 2010). A balance between the entrance and the exit of cations to preserve their concentrations at nutrient levels maintained by the cells as a response. Transport systems involved in cell homeostasis must be pivotal for the virulence of intracellular pathogens, such as *Mtb*. The large number of P-type ATPases encoded in the genome of *Mtb* is twice the size of those encoded in the genomes of saprophyte mycobacteria, such as *M. smegmatis*; this may strongly suggest the importance of this type of metal cation transporters to the virulence of the tubercle bacillus (Ward et al., 2011) *Mtb* P-type ATPases and related human counterparts. Albeit, *ctpH* gene having SERCA Ca ATPase as the human counterpart(Novoa-Aponte & Soto Ospina, 2014), they are divergent in comparison; thus, being prone to inhibition without affecting the host cells. Could a change in the frame of this gene favour *Mycobacterium Tuberculosis,* from being inhibited by the host cell due to its divergence in the human counterpart.

Therefore more *invivo* laboratory studies need to be done to investigate the effect of having both a frameshift and missense mutation in the *ctpH* gene function and also instabilities in the Uganda I genotype genome.

## Methods and materials

### Data availability

Publicly available whole genome sequences of Mycobacterium tuberculosis were downloaded from the sequence read archive, which is housed under NCBI. This was done using the Sequence Read Archive toolkit (http://ncbi.github.io/sra-tools/). The reads were then checked for quality using FastQc software package (http://www.bioinformatics.babraham.ac.uk/projects/fastqc/). All the reads that did not meet the threshold quality scores were trimmed off using trimmomatic (Bolger, A. M., Lohse, M., & Usadel, B. (2014)). Variant calling was then performed using snippy 4.6.0 (https://github.com/tseemann/snippy). Snippy requires a reference genome and raw reads as input. For our analysis, we used Mycobacterium tuberculosis H37Rv complete genome as the reference. This was downloaded from NCBI as a full genbank file and the accession number used is AL123456.

### Bioinformatics analysis

#### Mapping and variant calling

FastQ reads were trimed using Trimmomatic v 0.33 478 (SLIDING WINDOW: 5:20) (50) to remove low quality reads shorter than 20 bp for the downstream analysis. Variant calling was then performed using snippy 4.6.0 (https://github.com/tseemann/snippy). Snippy first did a reference based mapping using bwa (Li H. and Durbin R. (2009) Fast and accurate short read alignment with Burrows-Wheeler Transform. Bioinformatics, 25:1754-60.‘ [PMID: 19451168]) with the M. tuberculosis H37Rv as the reference genome. The reference genome was downloaded from NCBI as a full genbank file and the accession number used is AL123456.3. Followed was variant calling using freebayes (Garrison E, Marth G. Haplotype-based variant detection from short-read sequencing. *arXiv preprint arXiv:1207.3907 [q-bio.GN]* 2012) and then annotation of variants using snpEff (A program for annotating and predicting the effects of single nucleotide polymorphisms, SnpEff: SNPs in the genome of Drosophila melanogaster strain w1118; iso-2; iso-3.”, Cingolani P, Platts A, Wang le L, Coon M, Nguyen T, Wang L, Land SJ, Lu X, Ruden DM. Fly (Austin). 2012 Apr-Jun;6(2):80-92. PMID: 22728672 [PubMed - in process]). Using customized scripts, isolates were divided into Ugandan and non ugandan. Ugandan for those that had the Ugandan genotype genetic marker. The Ugandan isolates were then divided into Uganda I and II using their respective genetic marker. Those that didn’t fall in either of the Uganda sub lineages I or II were regarded as notag.

#### Mixed infection analysis

The NOTAG genomes were further analysed for mixed infection using TBProfiler v2.8.12 (https://genomemedicine.biomedcentral.com/articles/10.1186/s13073-019-0650-x). TB-Profiler is a software which allows users to analyse *M. tuberculosis* whole genome sequencing data to predict lineage and drug resistance. The software searches for small variants and big deletions associated with drug resistance. It will also report the lineage. By default it uses Trimmomatic to trim the reads, BWA (or minimap2 for nanopore) to align to the reference genome and GATK (open source v4) to call variants. Those genomes that flanked as mixed infections were excluded in the downstream analysis.

#### Phylogenetic construction

All the Ugandan genomes were subjected to multiple core SNP site alignment using snippy-core. The multiple sequence alignment was then used to infer phylogeny using ete3 (*ETE 3: Reconstruction, analysis and visualization of phylogenomic data.* Jaime Huerta-Cepas, Francois Serra and Peer Bork. Mol Biol Evol 2016; doi: 10.1093/molbev/msw046) build based on the full model test bootstrap workflow. ETE is a Python programming toolkit that assists in the automated manipulation, analysis and visualization of phylogenetic trees. To differentiate the Uganda I, Uganda II and notags, we dadded a suffix of Uganda I, Uganda II or notag depending on their identity before running the snippy-core. Snippy-core produced a core.aln file which contained the alignment for core SNPs. We then inferred a phylogenetic tree using ete3 with application of full model test bootstrap. The topology was annotated and coloured using iTOL v5 (Letunic I and Bork P (2019) Nucleic Acids Res **doi: 10.1093/nar/gkz239** *Interactive Tree Of Life (iTOL) v4: recent updates and new developments*)

#### Pan-genome analysis

SPAdes genome assembler v3.14.1(http://bioinf.spbau.ru/spades) denovo assembled the remaining Uganda genotype genomes. The outputs from SPAdes were checked for quality using QUAST Alexey Gurevich, Vladislav Saveliev, Nikolay Vyahhi and Glenn Tesler,QUAST: quality assessment tool for genome assemblies,*Bioinformatics*(2013)29(8):1072-1075.doi:1 0.1093/bioinformatics/btt086First published online: February 19, 2013. Scaffolds were then annotated by prokka v1.4.6 and the pan genome calculated by roary v3.11.2. Pan genome wide association was performed using scoary v1.6.16. SPAdes is a de Bruijn graph-based assembler, which is noteworthy for its approach in applying multiple de Bruijn graphs (each built with different k-mer sizes) to better handle the large variations in coverage across the genome that are a characteristic of single cell sequencing, as well as a novel method for handling paired end information. It begins its assembly process by using multisized de Bruijn graphs for constructing the assembly graph while detecting and removing chimeric reads. Next, distances between the k-mers are estimated for mapping the edges of the assembly graph. Afterwards, a paired assembly graph is constructed and SPAdes outputs a set of contiguous DNA sequences (contigs). Prokka is a software tool for the rapid annotation of prokaryotic genomes. A typical 4Mbp genome can be fully annotated in less than 20 minutes. Roary(https://sanger-pathogens.github.io/Roary/) is a high speed standalone pan genome pipeline, which takes annotated assemblies in GFF3 format (produced by Prokka (Seemann, 2014)) and calculates the pan genome. Scoary (https://github.com/AdmiralenOla/Scoary) is a software designed to take the gene_presence_absence.csv file from roary as well as a traits file created by the user and calculate the associations between all genes in the accessory genome and the traits. It reports a list of genes sorted by strength of association per trait. Scoary outputs a single csv file per trait in the traits file. It uses comma “,” as a delimiter. The results consist of genes that were found to be associated with the trait, sorted according to significance. (By default, Scoary reports all genes with a naive p-value < 0.05, but the user can change the cut-off value and use adjusted p-values instead), their corresponding sensitivity and specificity. Scoary also provides a list containing a number of trait-positive isolates this gene was found in or absent in and also lists where the number of trait-negative isolates this gene was found in or not.

## Abbreviations

TB: Tuberculosis
MTBC: Mycobacterium Tuberculosis Complex
Mtb: Mycobacterium Tuberculosis
MTB: Mycobacterium Tuberculosis
EPTB: Extrapulmonary Tuberculosis
PTB: Pulmonary Tuberculosis
SNP: Single Nucleotide Polymorphism
RD: Regions of Difference
MIRU-VNTR: Interspersed repeat units - variable number of tandem repeats
RFLP: Restriction, Fragment Length Polymorphism
ATP: Adenosine Triphosphate

## Acknowledgements

We acknowledge Gerald Mboowa and Willy Ssengooba for their kind support throughout this project.

PN is a MUII-Plus bioinformatics strategic internee. MUII-Plus is funded through the DELTAS Africa Initiative grant. The DELTAS Africa Initiative is an independent funding scheme of the African Academy of Sciences (AAS)’s Alliance for Accelerating Excellence in Science in Africa (AESA) and supported by the New Partnership for Africa’s Development Planning and Coordinating Agency (NEPAD Agency) with funding from the Wellcome Trust (WT) grant #107742/Z/15/Z and the UK government. The views expressed in this publication are those of the author (s) and not necessarily those of AAS, NEPAD Agency, Wellcome Trust or the UK government. The funders had no role in study design, data collection and analysis, decision to publish, or preparation of the manuscript.

